# Microbial abundance estimation and metagenomic studies of holcomb creosote superfund site soil sample

**DOI:** 10.1101/2024.03.19.585654

**Authors:** D.S Kishore, C.N Prasantha, Majji Rambabu, Karthick Vasudevann, K.R. Dasegowda

**Affiliations:** Department of Biotechnology, School of Applied Science, REVA University

**Keywords:** METAGENOMICS, MG-RAST, NGS, MEGA

## Abstract

The use of genomic sequencing has greatly improved our ability to profile the microbial communities associated with the environment and host. Among the most common applications of metagenomics is assessing microbiome biodiversity. Holcomb Creosote Superfund Site Soil is located in the North Carolina area, 16s metagenomic Sequencing was carried out from the soil of Holcomb Creosote Superfund Site. To describe the taxonomic diversity and functional profiles of this environment, metagenomic DNA sequence was extracted from a metagenomics library generated from the Holcomb Creosote Superfund Site Soil by 16s metagenomic Sequencing. The DNA was shotgun-sequenced using Illumina and analysed using the MG-RAST server. A soil sample from a large metagenomes sequence collection acquired from shotgun sequencing was investigated for Duke University. Overall sequences in the dataset were 1,34,00,509 comprising a total read length of 4,03,05,03,377 base pairs. The categorization of 8 species was based on the analysis of taxonomic data. The metagenome sequence was submitted by Duke University, Alexander McCumber to NCBI database and can be accessed with the SRA accession number SRX8095153. An online metagenome server (MG RAST) using the subsystem database revealed bacteria had the highest diversity profile revealed that the most abundant domain was 92.3% of bacteria, 5.6% of Eukaryota, 0.1 % of viruses, 1.4% of archaea, and 0.6% of unclassified sequences.The most abundant were Firmicutes (20.5%), and Proteobacteria (10.3%) followed by Actinobacteria (18.4%) and Acidobacteria (7.5%). The functional profile showed an abundance of genes related to subsystems (16.9%), carbohydrates (19.5%), cell wall and capsule (6%), miscellaneous (8.8%), protein metabolism (10%), amino acids and derivatives (14.7%), DNA metabolism (3.4%), cofactors, vitamins, prosthetic groups, pigments (9.2%), membrane transport (1.9%), RNA metabolism (4.4%) and fatty acids, lipids, and isoprenoids (3.6%). This dataset is useful in bioprospecting studies with application in biomedical sciences, biotechnology and microbial, population, and applied ecology fields.

## INTRODUCTION

Soil is the most diverse natural environment on Earth. The soil microbial communities harbour thousands of different prokaryotic organisms, one gram of soil contain between 2,000 to 18,000 separate genomes worth of very wide genetic information. One of the most crucial concerns in the study of soil ecology is to uncover the complex relationships between microbial compositions and functional diversity in soil (Xu et al., 2014). Although soil microbial communities have been studied for many years, recent advances in high-throughput molecular techniques have enabled microbial ecologists to characterise the taxonomic, phylogenetic, and functional diversity of soil microbial communities to an extent that was previously unthinkable. (Fierer et al. 2012).

A growing number of researchers are using genomics for functional analyses of the complete genome. Transcripts of whole genome RNA (transcriptomic), proteins (proteomics), and metabolites (metabolomics) are examples of functional analytical features. These fields of research can help us better understand the structure and function of biological systems (Chiriac & Murariu et al. 2021). In applied soil science, shotgun metagenomics techniques have the benefit of amplifying sequences from throughout the genome, including coding sections, resulting in a high sequencing depth but at a greater cost. 16S Meta barcoding is a diagnostic technique for bacterial and archaeal identification used in clinical sciences and plant ecology (Senn et al. 2022). Targeted 16S rRNA amplicons sequenced by a variety of methods are currently being used to examine the microbiota composition in detail. This method is quicker and less expensive than shotgun metagenomics (Andrade et al., 2020; Koringa et al., 2019; Li et al., 2016).

Metagenomics has grown in importance as a driving factor for discoveries in microbial ecology and biotechnology, as well as a crucial approach for investigating the microbial universe, since its first use in soil. Our capacity to assess microbial diversity and functional potential in every environment expanded tremendously as sequencing technology got cheaper, quicker, easier to use, and had greater throughput. The availability and capacity of computational approaches are critical for successful metagenomics applications. We emphasize the new information gained through metagenomics applications in Earth’s many ecosystems, as well as the analytical techniques that allow such findings (Taş et al.2021).

As the cost of sequencing falls and more researchers create more and larger datasets, the demand for economic analysis rises. Several visualizations have been added to the MG-RAST (Meta Genome Rapid Annotation using Subsystems Technology) suite of tools to let people study datasets via a simple web-based platform (Antonopoulos et al., 2010). MG-RAST is one such platform that processes hundreds of submissions every day, accumulating more than 0.5 terabytes in a single 24hour period (Meyer et al., 2018). Other repositories include the MG-RAST server (Glass et al., 2011) and microbe (Hurwitz, 2014) MG-RAST at the phylum- and family level (Kaszubinski et al., 2019) as well as compositions at the genus-level (Plummer et al., 2015). The objective of the study of metagenomics analyses is to study the abundance, taxonomic prediction, species richness, species diversity and functional diversity were analyzed.

## Materials and Methods Data

### Description

The dataset comprises raw Illumina paired-end reads obtained from shotgun metagenomics sequence of DNA extracted from a 16s metagenomic study performed on polyaromatic hydrocarbon and heavy metal polluted soils from the Holcomb Creosote Superfund site using Illumina Miseq.

There are 4,03,05,03,377 sequences in the raw sequencing data 1.19 M with a mean sequence length of 220 ± 8 bp. The FASTQ format data read files submitted to the NCBI database as part of the study BioProject No. PRJNA385836 (https://www.ncbi.nlm.nih.gov/bioproject/PRJNA385836)BioSample accession number SAMN14573447 (https://www.ncbi.nlm.nih.gov/biosample/SAMN14573447) and SRA accession number: SRX8095153 (https://www.ncbi.nlm.nih.gov/sra/SRX8095153) was selected as data set for metagenomic studies. The sequences quality check using MG-RAST analysis. Data at the phylum level, a rarefaction curve, and α-diversity are all included in the dataset.

Furthermore, the dataset includes the distribution. The dataset also includes the distribution of possible functional categories for COGs, KOs, NOGs, and SUBSYSTEMs at the greatest level supported by these functional hierarchies.

### Required data for MG-RAST

In today’s big data bioinformatics, the barrier to data and result reuse is a serious concern. It may be difficult to compare the results of an expensive computational approach to those of a different laboratory if the processes used aren’t equal (potentially compromising the integrity of the study). Another frequent strategy is to avoid reusing previous results and instead do an expensive reanalysis of both data sets, thus replicating the previous effort. Because no reviewer can be expected to retrace the scientists’ steps while repeating their computer labour, data-driven research can no longer be peer-reviewed. If data and results (including interim discoveries) were presented as repeatable entities, the problem of data uncertainty and costly recompilations would be removed. MG-RAST created a web page to view, analyzed, and download results so that they can be used for comparison with other tools (Kibegwa et al., 2020).

### Sequence data supported

Currently, the MG-RAST system accepts shotgun and amplicon metagenomes, as well as metatranscriptomes, in FASTQ (preferred) or FASTA format, from any platform. Users who want to provide assembled or pre-analyzed datasets are encouraged to additionally submit the ‘raw’ data, which has not been edited. For submission to MG-RAST, a minimum read length of 75 base pairs is required, as well as a minimum dataset size of 1 megabase. (Wilke et al., 2016)

### Updates to metadata

MG-RAST employs spreadsheets based on GSC metadata standards to manage metadata. With Metazen, the system validates metadata spreadsheets automatically. The Genomic Standards Consortium (GSC) has created metadata standards for genomic, metagenomic, and amplicon (e.g., 16S rRNA) sequence collections that are generally recognized (Bischof et al., 2014).

## RESULT

Many studies have made use of the publicly accessible suite of metagenomic analysis tools known as MG-RAST. To predict bacteria and viruses associated with respiratory disease in the current experiment, the MG-RAST server was employed to assist in the analysis of virome datasets at the microbiological level. It supports the analysis of raw reads, fna, or fast files in addition to contigs, providing higher-quality data for more precise analysis. The datasets uploaded to MGRAST contained 1,34,00,509 sequences with a combined base pair count of 4,03,05,03,377. One of the first places to look for any data collection is the purpose and feature breakdown at the top of the overviewpage. The pie charts at the top of the overview page are a good place to start (Fig. 1(A)) categorize the provided sequences into multiple categories based on their QC results: failed (Bule), Predict feature (Yellow), and unknown (Green). (Fig. 2(B)) divides the anticipated characteristics into unknown protein (Green), annotated protein (Bule), and ribosomal RNA (Yellow).

**FIG. 1.**
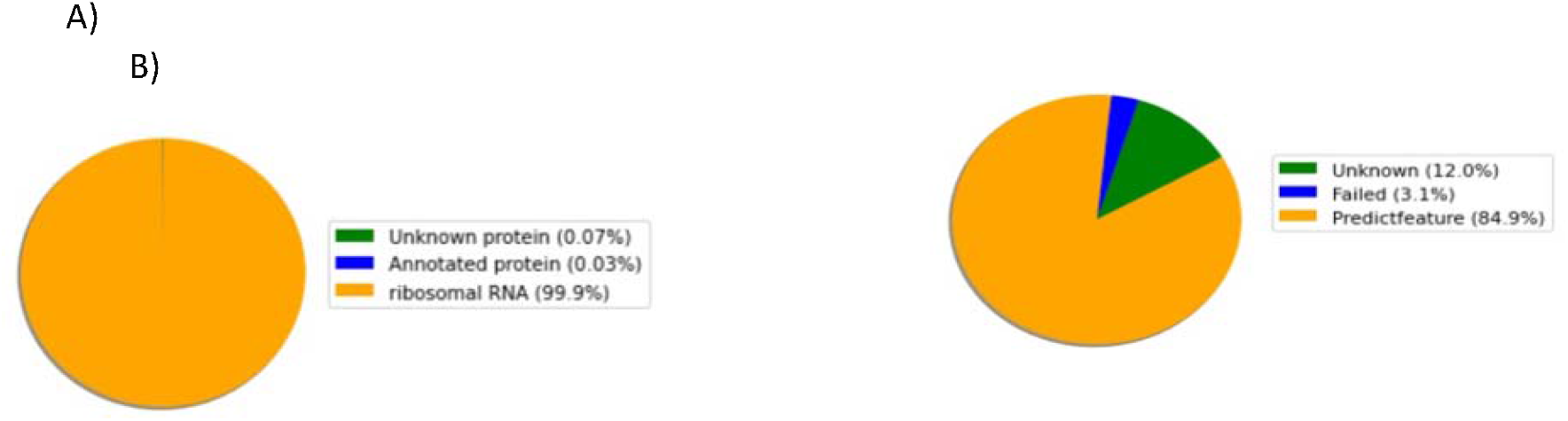
Overview of Provenance. (A) Predict feature, (B) Sequence feature.

**FIG. 2.**
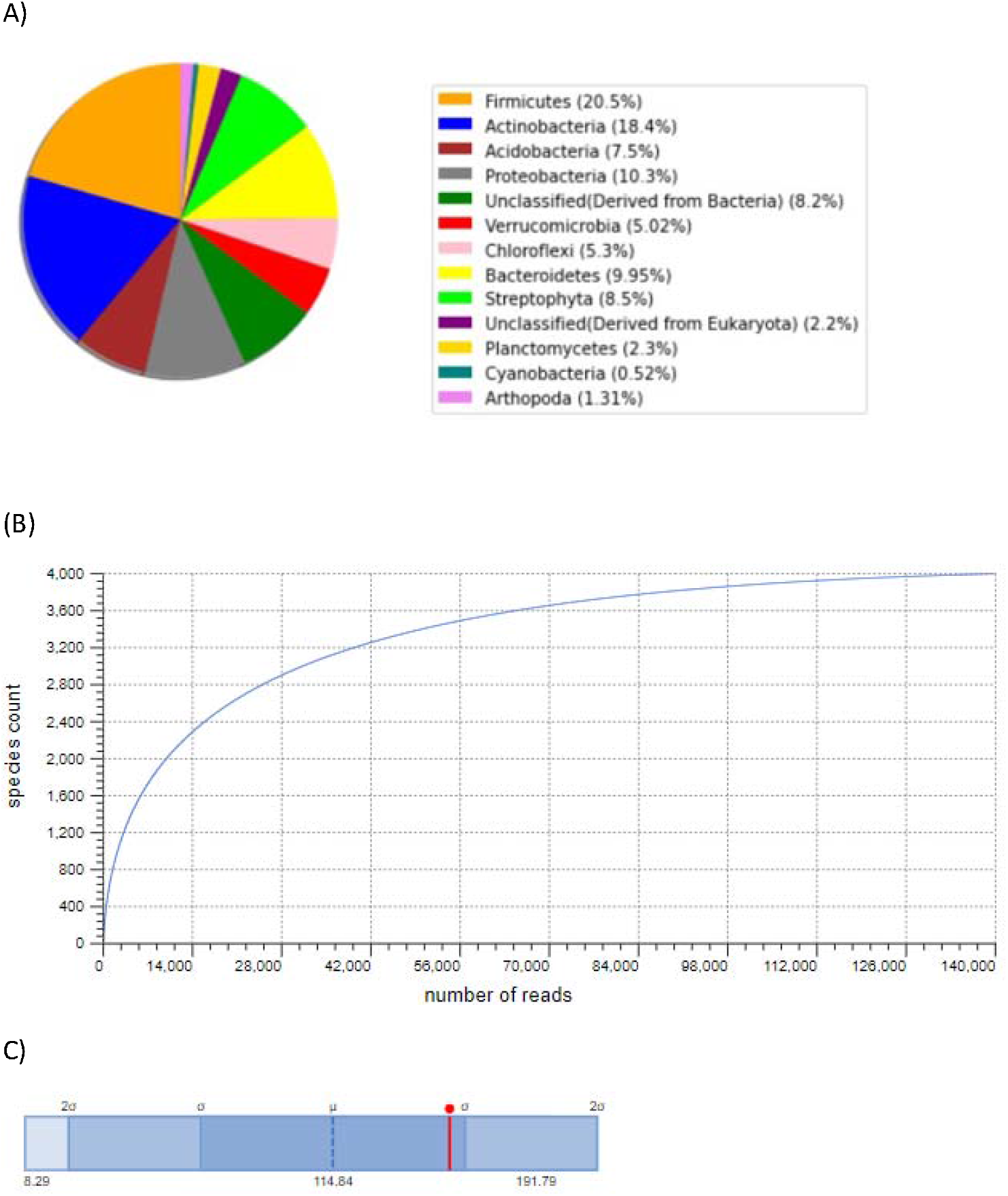
Overview of taxonomic hits distribution. (A) Phylum (B) Rarefaction curve (C) Alpha diversity. **FIG. 2. (A)** Representation of the phylum-level composition of the samples from MG-RAST had considerably greater relative abundances of unclassified and unidentified sequences, respectively. Protein similarities to entries in the “RefSeq” protein databases are used to represent the distribution of taxonomic groups at the phylum level in Figure 1. Taxonomic diversity of the soil 16s metagenomic sequencing from the Holcomb Creosote Superfund Site. The most prevalent domain identified using the RefSeq database resource in MG-RAST was Firmicutes (20.5%), Actinobacteria (18.4%), Acidobacteria (7.5%), Proteobacteria (10.3%), Unclassified (Derived from Bacteria) (8.2%), Verrucomicrobia (5.02%), Chloroflexi (5.3%), Bacteroidetes (9.95%), Streptophyta (8.5%), Unclassified (Derived from Eukaryota) (2.2%), Planctomycetes (2.3%), Cyanobacteria (0.52%), Arthropoda (1.31%) of sequences remained unannotated. **FIG. 2. (B)** Rarefaction plot representation with a species richness curve annotated (i.e., the number of unique species). Figure 2. This curve shows the overall number of various species annotations as a function of the number of sequences sampled. The rarefaction view can only be used to display taxonomic data. The rarefaction curve of annotated species richness is shown in a FIG. 2. (B) **FIG. 2. (C)** The range of -diversity values in the project the data collection is a part of are shown in the project’s alpha diversity graphic representation. The ranges of the standard deviations (and 2) are shown by different colours, and the lowest, maximum, and mean values are given. Red highlights indicate the diversity of this metagenome. In this part, we offer an assessment of the alpha diversity based on the taxonomic annotations for the predicted proteins. The alpha diversity is presented with other metagenomes in the same study Fig. 2. (C). The alpha diversity estimate, which is a single integer, summarises how many species-level annotations are present in a dataset. The Shannon diversity index is a logarithm of the relative abundances of annotated species, weighted by abundance. The Shannon diversity is used to calculate the species richness. Pi represents the percentages of annotations in each category of species, and 10 P I pi log is used to compute richness (pi). Shannon defines species richness as the “number of effective species”. The amount of annotations for each species is divided by the total number of annotations to create each p. We use all of the source databases that MG-RAST used to generate its species-level annotations.

**FIG. 3.**
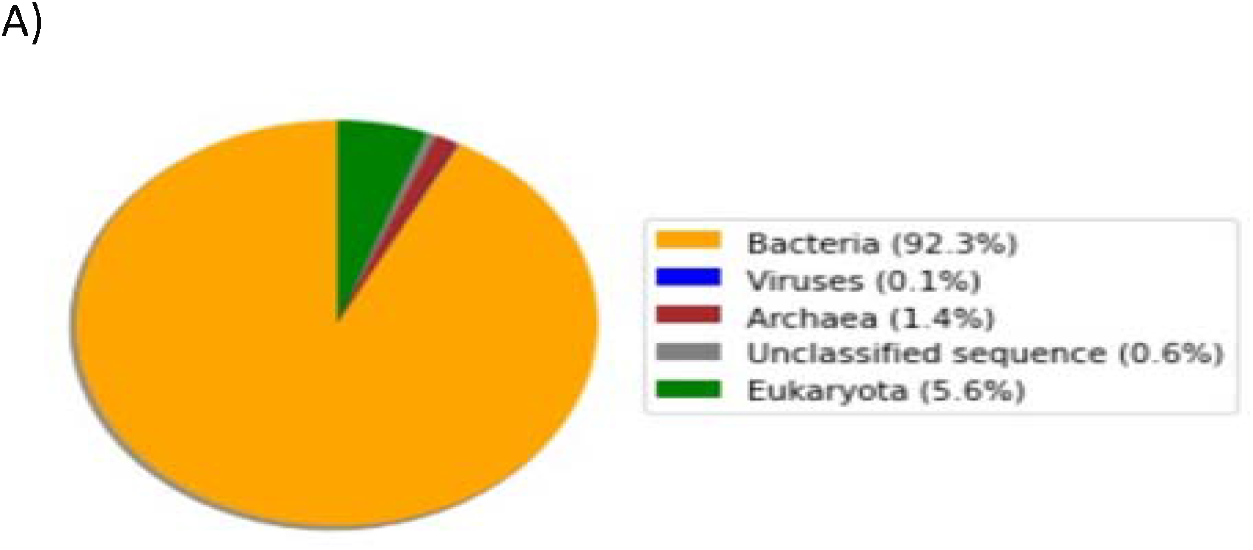
**(A)** Figure 3 shows the taxonomic composition of 30 samples as assessed by MG-RAST. The average relative abundance differences between Unclassified, Bacteria, and Eukaryota were statistically significant using MG-RAST, but not for Archaea. The main reasons for these discrepancies were the much higher proportion of unclassified sequences in MG-RAST and the inaccurate classification of 16S rRNA amplicon sequences into Eukaryota and Viruses. The Archaea: Bacteria ratio for each sample was calculated using the absolute counts of hits obtained from each pipeline, and the results were compared using a two-sided paired t-test. It was discovered that there was a significant difference between the pipelines, primarily because of the different abundances of bacteria. The distribution of taxonomic categories at domain levels using protein similarities to entries in the ‘RefSeq’ protein databases were bacteria (92.3%), eukaryote (5.6%), viruses (0.1%), archaea - (1.4%), and unclassified sequences (0.6) of sequences remained unannotated. **FIG. 3**. Overview of taxonomic hits distribution. (A) Domain.

**FIG. 5.**
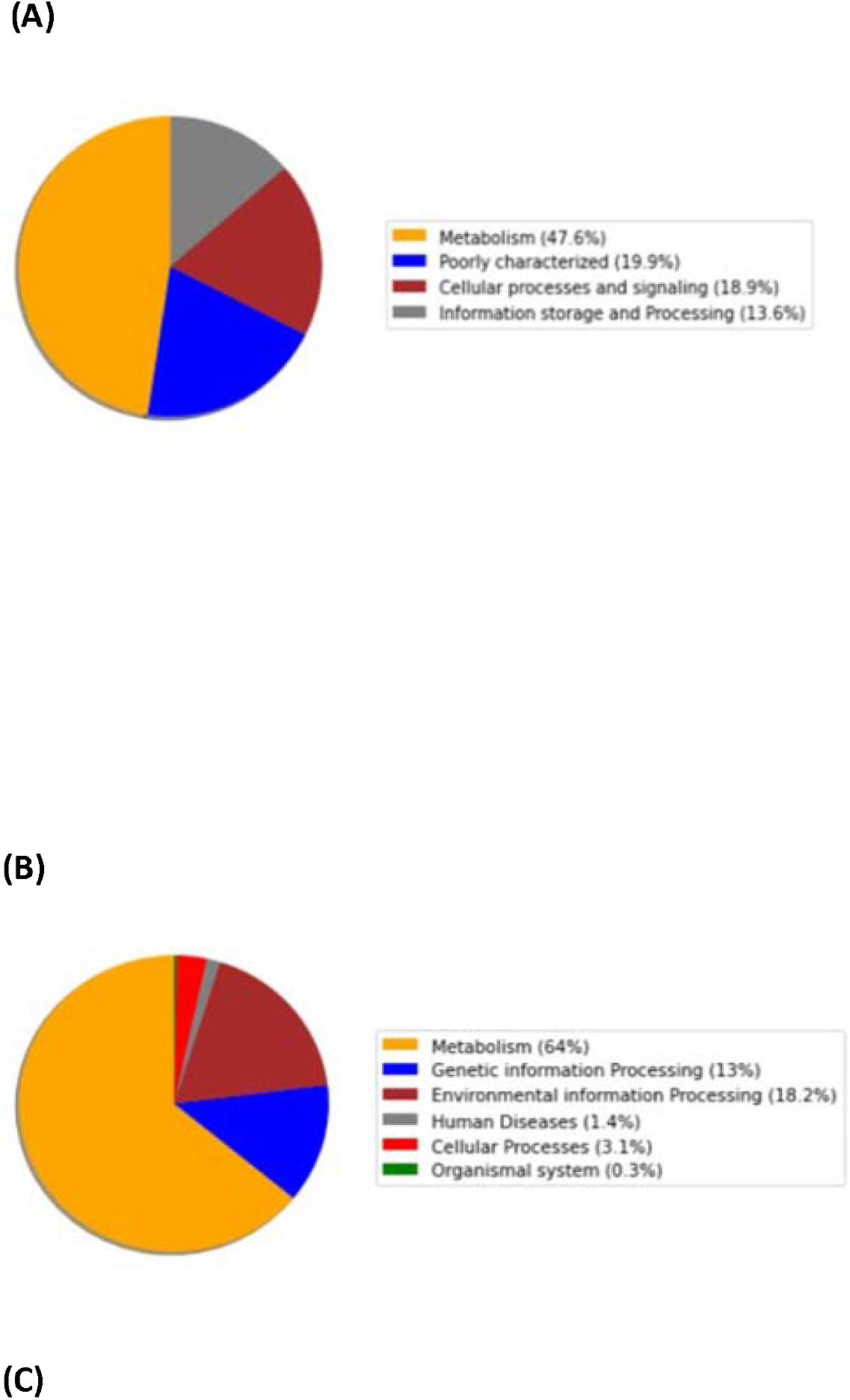
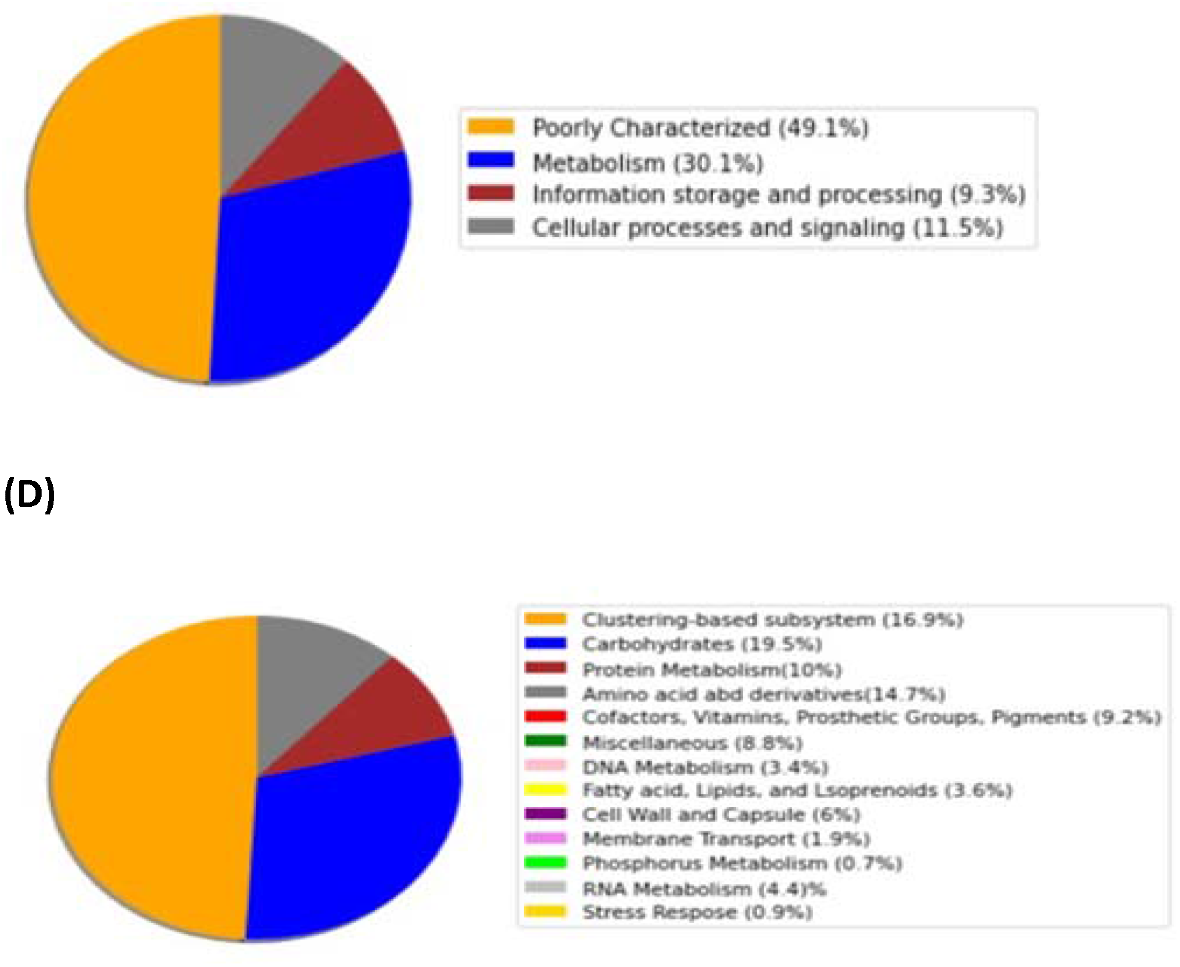
The neighbour-joining approach was used to root the phylogenetic tree based on 16S rDNA sequences. 0.0 substitutions per site, bar.

**FIG. 6.**
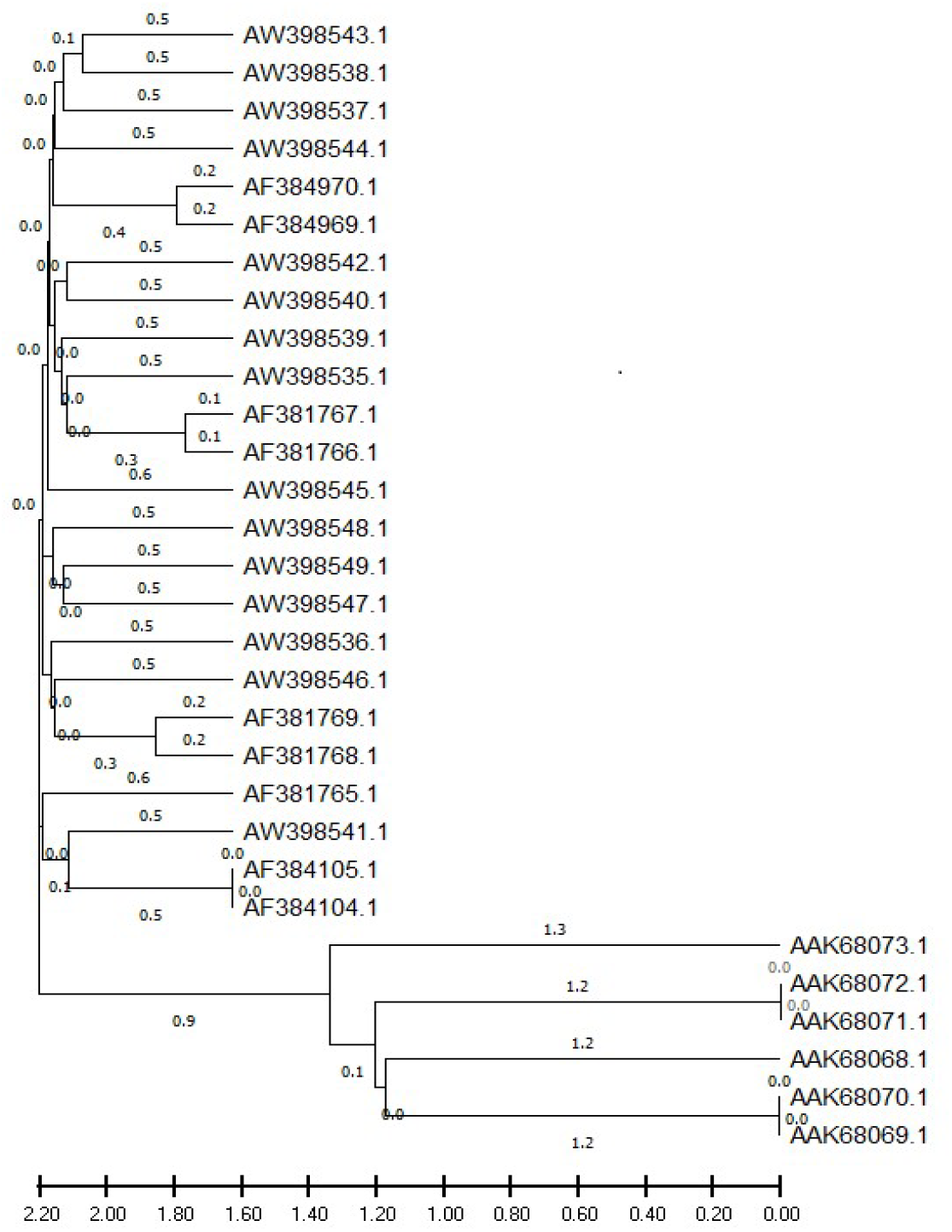
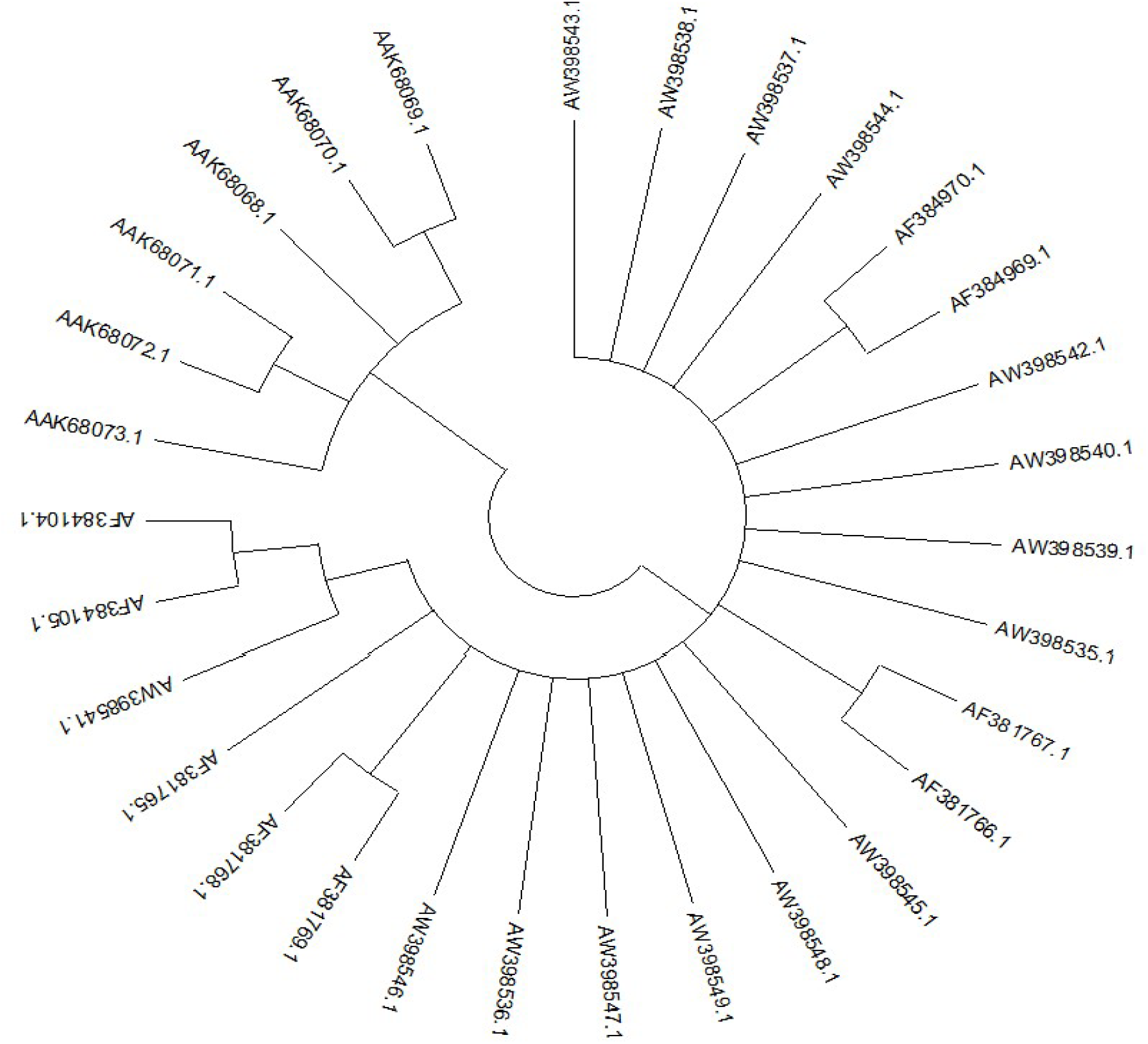
Phylogenetic tree built utilising the test organisms 16S rDNA sequences and appropriate reference or type strains collected from the nucleotide database is displayed. The tree topology created using the maximum likelihood approach supported the phylogenetic tree topology created using the neighbour-joining method. It was found that there were six clusters of operational taxonomic units (OTUs) that supported the internodes with bootstrap values more than 60%. By creating a phylogenetic tree, it is possible to characterise the evolutionary connections between the sequences under study. The sample sequence is located in a different clade than the sample (AF381765.1). The building of the phylogenetic tree between the samples revealed a trustworthy bootstrap value that was in the region of 50% bootstrap analysis with a value of more than 80% demonstrating the accuracy of the grouping. **FIG. 6**. The neighbour-joining phylogenetic tree built using the 16S rDNA sequences as the basis rooted the untree. 2.0 replacements per site, bar.

As a result of the functional categorization of sequences, 12.0% of predicted proteins had known functions, 84.90% of predicted proteins had unknown functions, and a minuscule 99.9% of ribosomal RNA genes had unknown functions.

The first pie charts classify the sequences submitted in this data set according to their QC results, and the 2nd breaks down the detected features into several categories.

## TAXONOMIC HITS DISTRIBUTION

Visualization of the phylum levels, rarefaction curve, and α-diversity of the microbiome from a composite soil sample taken from a16s using shotgun metagenomics using soils from the Holcomb Creosote Superfund site that were polluted with polychromatic hydrocarbons and heavy metals, metagenomic analysis was done utilising the Illumina Miseq platform. Using protein similarities to entries in the “RefSeq” protein databases, the distribution of taxonomic groupings at phylum levels.

## DOMAIN

Protein similarities to entries in the “refseq” protein databases are used to determine the distribution of taxonomic groupings at domain levels.

## SEQUENCE ANALYSIS

Raw Illumina paired-end reads were uploaded as FASTQ files, and the MG-RAST server version 4.0.3 was used for analysis. The COGs ontology, KOs ontology, NOGs ontology, and Subsystems ontology were used to indicate the functional categories after high-quality reads were submitted to analysis to predict, identify, and assign biological functions (gene annotations) to proteins and rRNA.

## FUNCTIONAL CATEGORY HITS DISTRIBUTION

Shotgun metagenomics of the microbiome from a composite soil sample taken from a 16s soil sample was conducted using the illuminate Miseq platform. In order to determine possible functional categories for COGs, KOs, NOGs, and SUBSYSTEMs, metagenomic analysis utilising the Illumina Miseq platform was carried out on polychromatic hydrocarbon and heavy metal-contaminated soils from the Holcomb Creosote Superfund site. Sequences for metabolism, information storage processing, cellular processing, and signalling were divided into categories by the functional analysis utilising COGs, KOs, NOGs, and SUBSYSTEMs. The sequences for poorly described (49.1%), metabolism (30.1%), information storage and processing (9.3%), cellular activities, and signalling were distributed both numerically and in terms of percentage in the subsystem technology (11.5). This exhibits the makeup of the microbial community, which aids in assessing the variety of the microorganisms.

## PHYLOGENETIC ANALYSIS

## Discussion

Metagenomics plays a crucial role in exploring microbial ecology and biotechnology, providing valuable insights into the microbial world. In a specific study, raw Illumina paired-end reads were obtained from shotgun metagenomic sequencing of soil samples contaminated with polyaromatic hydrocarbons and heavy metals. The data was analyzed using the MG-RAST system, which accepts metagenomic and metatranscriptomics data in FASTQ or FASTA format. The MG-RAST server facilitated the analysis of virome datasets, ensuring high-quality data for precise analysis.

The study focused on visualizing the phylum levels, rarefaction curve, and α-diversity of the microbiome using shotgun metagenomics of a composite soil sample. The analysis employed the Illumina Miseq platform, with the raw Illumina paired-end reads uploaded as FASTQ files. The MG-RAST server version 4.0.3 was utilized for the analysis.

The main objective of the metagenomic analysis was to identify functional categories such as COGs, KOs, NOGs, and SUBSYSTEMs. This analysis specifically targeted soil samples contaminated with polyaromatic hydrocarbons and heavy metals from the Holcomb Creosote Superfund site. The Illumina Miseq platform was used to generate the metagenomic data for this analysis.

## CONCLUSION

A soil sample was analyzed using MG-RAST, a metagenomics analysis tool, and it successfully identified microbial species associated with respiratory diseases. The analysis also led to the discovery of previously unknown species. The use of different data analysis tools with varying annotation databases results in different capabilities and workflows. The respiratory infection prevention techniques found in the soil sample were made possible by the enhanced sequence analysis provided by MG-RAST. The annotation process of MG-RAST facilitated the analysis of sequences and made it easier to interpret the results. The MEGA program allows for interactive and comparative phylogenetic analysis of various datasets, complementing the findings from MG-RAST.

